# Assessment of SARS-CoV-2 Specific CD4(+) and CD8 (+) T Cell Responses Using MHC Class I and II Tetramers

**DOI:** 10.1101/2020.07.08.194209

**Authors:** Yuri Poluektov, Pirouz Daftarian, Marc C. Delcommenne

## Abstract

The success of SARS-CoV-2 (CoV-2) vaccines is measured by their ability to mount immune memory responses that are long-lasting. To achieve this goal, it is important to identify surrogates of immune protection, namely, CoV-2 MHC Class I and II immunodominant pieces/epitopes and methodologies to measure them. Here, we present results of flow cytometry-based MHC Class I and II QuickSwitch™ platforms for assessing SARS-CoV-2 peptide binding affinities to various human alleles as well as the H-2 Kb mouse allele. Multiple SARS-CoV-2 potential MHC binders were screened and validated by QuickSwitch testing. While several predicted peptides with acceptable theoretical Kd showed poor MHC occupancies, fourteen MHC class II and a few MHC class I peptides showed promiscuity in that they bind with multiple MHC molecule types. With the peptide exchange generated MHC tetramers, scientists can assess CD4+ and CD8+ immune responses to these different MHC/peptide complexes. Results obtained with several SARS-CoV-2 MHC class I and II peptides are included and discussed.

## Introduction

While various antiviral drugs or passive antibody therapies are attractive approaches as a bridge to a vaccine, the consensus is that immunization and mounting durable immune memory against the SARS-CoV-2 is an unavoidable task facing today’s scientific community [1]. The basic research to uncover the exact mechanisms of how the SARS-CoV-2 virus is perceived and countered by the human immune system have been more difficult. In the effort to fight this outbreak, a greater understanding of how the viral proteins and their processed peptides stimulate the immune responses mediated by CD8+ and CD4+ T lymphocytes is needed. While some tremendous research has been undertaken with an impressive swiftness to determine the specific viral peptides that lead to an effective T cell response, many of the identified SARS-CoV-2-reactive T-cells were also found in healthy donors [2,3]. This, in turn, raises even more questions as to the nature of the viral peptides needed to mount an effective immune response. In addition, this creates a vital need for a fast and effective way to generate various MHC tetramers needed to detect the numerous specific T cells that will be researched while trying to determine the best possible viral peptides that would stimulate the most effective T cells.

The ability of a peptide to stimulate the immune system stems from such properties as the availability of the right T cell receptor, antigen processability by the antigen processing system that could generate the peptide in question, and the peptide’s ability to occupy the cleft of the MHC molecule [4,5]. It has been observed that more than 80% of MHC class I-bound peptides derived from a virus can be immunogenic [6]. At least in the case of viral peptides, peptides that that have a high binding affinity to MHC Class I molecules tend to also be immunogenic [7,8]. To this end, we have come up with an antigen presentation platform based on the QuickSwitch peptide exchange principle. This platform allows users to quantitate the relative ability of MHC molecules to bind a particular peptide, as well as to generate an MHC tetramer that can be used to stain T cells in Flow Cytometry experiments. This system relies on an irrelevant and weak binding peptide, called exiting peptide, which comes pre-loaded into the recombinant MHC molecule that you wish to study. When the QuickSwitch MHC molecule is presented with a competing peptide, the exiting peptide will be replaced proportionally to the MHC binding affinity of the competing peptide. In addition, a FITC-labeled antibody against the exiting peptide comes included in every kit so that the extent of the peptide exchange can be measured using a simple Flow Cytometry procedure with magnetic beads specific for the MHC molecule being tested. In this short study we would like to report how the QuickSwitch platform (MBL International) can be used to screen SARS-CoV-2 peptides for their ability to bind various MHC molecules. This short collection of findings helps avoid unnecessary experiments with SARS-CoV-2 peptides that cannot be presented by MHC molecules and allows for the construction of working MHC tetramers that can be used for productive CD8+ and CD4+ cell staining.

## Materials and Methods

The peptide binding tests to determine the peptide’s affinity for a specific MHC haplotype were performed on a 50 µg/mL (MHC concentration) solution of Tetramers for the MHC Class I molecules or a 100 µg/mL solution of recombinant MHC biotinylated molecules for MHC Class II. While it is possible to get valid peptide exchange results by decreasing the Tetramer or monomer concentration even further, it is not recommended, as some MHC haplotypes are much more sensitive to dilution than others generating different results depending on the MHC concentration.

### MHC Class I peptide exchange procedures

50 µL of the Tetramer at 50 µg/mL was mixed with 1 µL of Peptide stock diluted to 1 mM in H_2_O and 1 µL of the proprietary Peptide Exchange Factor corresponding to the particular MHC haplotype. HLA-A*02:01, HLA-A*24:02 and H-2 Kb received 1 µL of Peptide Exchange Factor #1 (PEF#1). HLA-A*11:01 received 1 µL of Peptide Exchange Factor #2 (PEF#2). And HLA-A*03:01 received 1 µL of Peptide Exchange Factor #3 (PEF#3). The resulting solution, containing 50 µg/mL Tetramer, 20 µM peptide to be loaded into the MHC molecule and 1 µL of the Peptide Exchange Factor was incubated in a conical-bottom 96 well PCR plate for 4 hours at room temperature (≈ 22°C).

### Capture assay to determine the binding affinity of a peptide for a particular MHC haplotype

Following the incubation, the percentage of MHC molecules that were loaded with the test peptide from the total MHC molecules containing the exiting peptide was determined by performing a capture assay described in the MBL International QuickSwitch™ Quant Tetramer Kit Data Sheet. Briefly, 20 µL of magnetic beads linked with an anti HLA-ABC antibody (provided in the kit) were added into sample wells of a V-bottom 96-well microtiter plate with 5 µL of the Tetramer peptide exchange solution (from the 52 µL total volume). The microplate was shaken for 45 minutes at 550 RPM, protected from light with a sheet of aluminum foil. Following the shaking incubation, all sample wells were washed with 150 µL of Assay Buffer (provided in the kit, must be diluted 10x before use), utilizing the magnetic qualities of the beads by letting them sediment while the plate is immobilized on a magnetic tray. The excess buffer was decanted out of the plate by flicking while it was being held tight to the magnetic tray. After rinse each well received 25 µL of Exiting Peptide Antibody-FITC solution (provided in the kit, must be diluted 25x before use). The microplate containing the magnetic beads with the bound MHC tetramers and the exiting peptide antibody was shaken again for 45 minutes at 550 RPM, protected from light with a sheet of aluminum foil. The samples were washed with another 150 µL volume of Assay Buffer and resuspended in 150 µL of Assay Buffer before the beads read on the flow cytometer.

### MHC Class II peptide exchange procedures

For MHC Class II peptide exchange we followed the parameters of QuickSwitch Quant HLA-DRB1*01:01, HLA-DRB1*04:01 and HLA-DRB1*15:01 assays. 25 µL of a 100 µg/mL biotinylated MHC Class II QuickSwitch molecule was mixed with 3 µL of Peptide stock diluted to 10 mM in 100% DMSO and 2 µL of Emulsifier #1 (provided in the kit). The resulting solution, containing 100 µg/mL MHC Class II molecule, 1 mM peptide (or tenfold dilutions) to be loaded into the MHC Class II molecule and 2 µL of the proprietary Emulsifier #1 was incubated in a conical-bottom 96 well PCR plate overnight (>10 hours) at 37°C.

The capture assay was performed with nearly identical procedures to the ones used for the MHC Class I Tetramers. 25 µL of streptavidin conjugated magnetic beads (provided in the kit) were used to capture 1 µL of the 30 µL MHC Class II peptide exchange reaction. The MHC molecules were captured by the magnetic beads after shaking the V-bottom 96-well microtiter plate for 45 minutes at 550 RPM. Following a 150 µL wash with Peptide Exchange Assay Buffer #2 (provided in the kit, must be diluted 10x before use) the exchanged MHC Class II molecules were stained with 25 µL of Exiting Peptide Antibody-FITC (provided in the kit). The antibody was allowed to stain the captured MHC Class II molecules by shaking the plate for 45 minutes at 550 RPM. The beads were then washed with another 150 µL of Peptide Exchange Assay Buffer #2 and resuspended in 150 µL of assay buffer.

### Flow Cytometry

The experiments were performed on the CytoFLEX LX (Beckman Coulter) flow cytometry instrument. The FSC and SSC voltages, gains and thresholds were adjusted so that the magnetic beads used for the capture assay were detected as a unique population separate from the background signals. The flow cytometer was then used to measure the Median Fluorescence Intensity of the FITC signal that resulted from the Exiting Peptide Antibody binding to the captured MHC tetramers on the anti HLA-ABC antibody covalently linked beads.

### Peptides

All peptides for this experiment were synthesized at the 95% purity level. The peptides were diluted to 10 mM stocks in 100 % DMSO and stored at −20°C. Further peptide dilutions before each binding experiment were done with HPLC grade H_2_O.

### In Silico prediction of peptide binding affinities for MHC molecules

All of the theoretical binding affinities were derived from the Immune Epitope Database (IEDB) web resource funded by NIAID. This website catalogues experimental data on antibody and T cell epitopes resulting from numerous studies in the context of infectious disease, allergy, autoimmunity and transplantation. The accumulated data is then compiled and analyzed to allow the prediction of peptide binding affinities for specific MHC molecules.

## Results

The ability of a number of SARS-CoV-2 virus derived peptides as potential binders to MHC molecules were tested. A few specific SARS-CoV-2 peptides with a high theoretical binding affinity for the MHC molecules in question, as determined by the Immune Epitope Database (IEDB) web resource were selected from a large list of MHC potential binders (supplemental tables 3 and 6). Those peptides were then loaded into MHC molecules using the parameters of the QuickSwitch kits.

In silico prediction, while an indispensable tool for modern studies on T cell stimulation, does not always predict the right peptide that would be able to bind to the MHC molecule’s peptide binding groove, not to mention, stimulate the T cells needed to fight infection or disease, especially when it comes to MHC Class II molecules [10]. The following results, obtained with recombinant MHC molecules, can be used in the future to determine the nature of the T cell immune responses to SARS-CoV-2 without questioning what viral peptide sequences can be presented by which MHC molecules. In addition, it will be simplify custom MHC tetramer generation with peptides that have shown to be good binders.

Using the HLA-A*02:01, HLA-A*03:01, HLA-A*24:02, HLA-A*11:01 and H-2 Kb QuickSwitch™ Quant Tetramer Kits, we were able to identify a number of peptides that bound to each MHC haplotype (Table 1). The identified SARS-CoV-2 peptides effectively displaced the exiting peptide which was selected for each individual MHC haplotype to easily dissociate from the MHC molecule to allow for the binding of even moderately strong peptides. When more than 75% of the exiting peptide is exchanged with a target peptide, the resulting tetramer can be used for CD8+ T cell staining. An exchange rate of higher than 90% corresponds to binding affinities in the low nanomolar range, while exchange rates lower than 65% are indicative of weak peptide-MHC interactions (data not shown). In contrast to MHC class II binding peptides there is very little binding promiscuity among class I alleles with one notable exception: HLA-A3 and HLA-A11. These 2 alleles are tightly related and all the tested peptides that were highly exchangeable on HLA-A3 also happened to be good HLA-A11 binders.

**Table 1:**
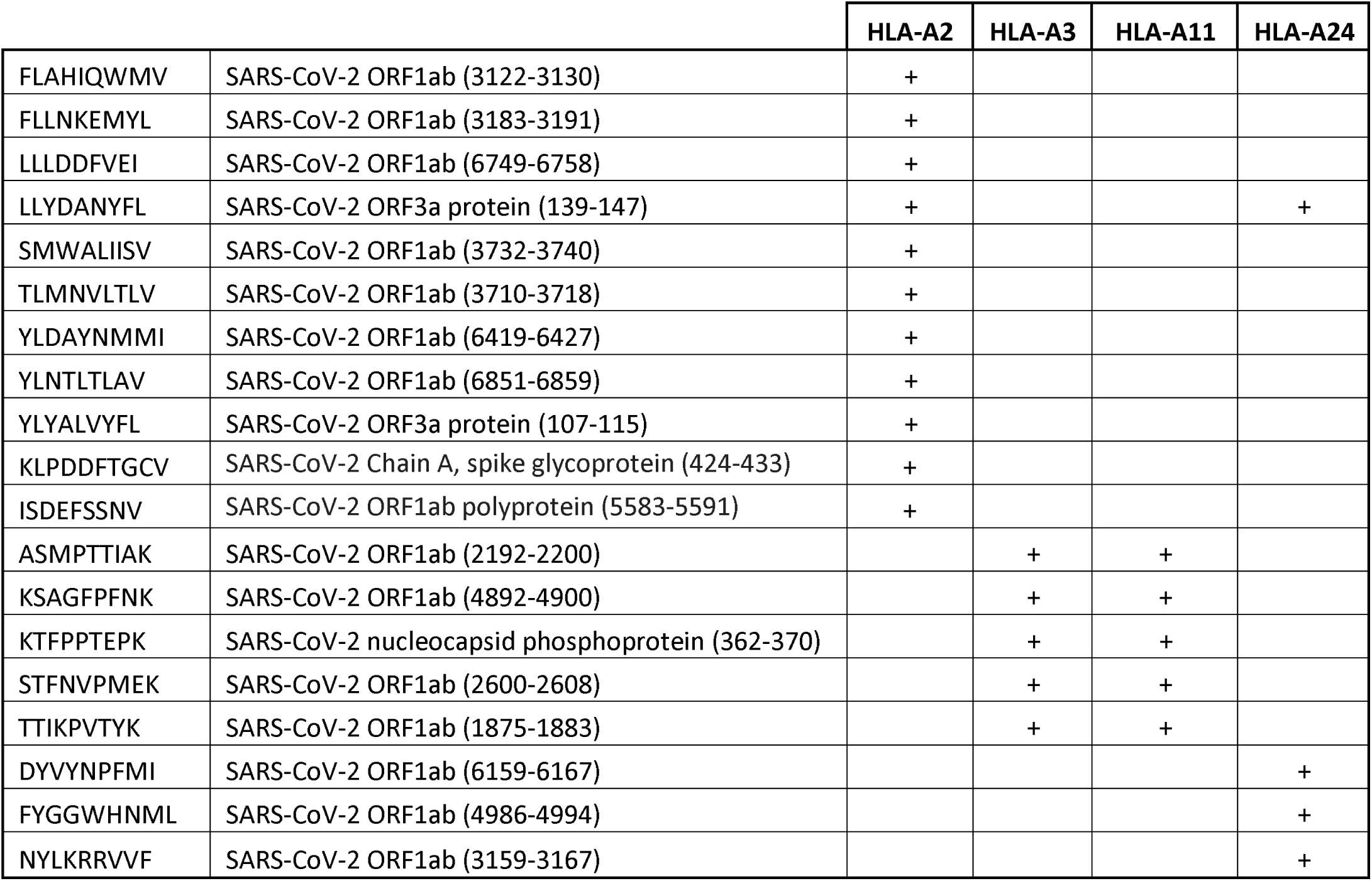
Summary of SARS-CoV-2 peptide binding to human MHC class I molecules. Peptides from different SARS-CoV-2 proteins were selected for strongest binding on the most frequent human MHC class I alleles: HLA-A2, A3, A11 and A24. All of the identified peptides were able to exchange at least 75% of the exiting peptide and could be used to make custom Tetramers for CD8+ T cell staining.

To address similar studies in mouse models, we attempted to assess the utility of the platform in a mouse MHC haplotype. Mouse H-2 Kb specific SARS-CoV-2 peptides which could be used for T cell immunomonitoring in mouse models of COVID-19 studies were identified, should they become available. A set of predicted peptides were tested and determined their peptide occupancies in the groove of H-2 Kb molecules.

Interestingly, two of the identified H-2 Kb peptides were also determined to be good HLA-A*02:01 binders (Table 2).

**Table 2:**
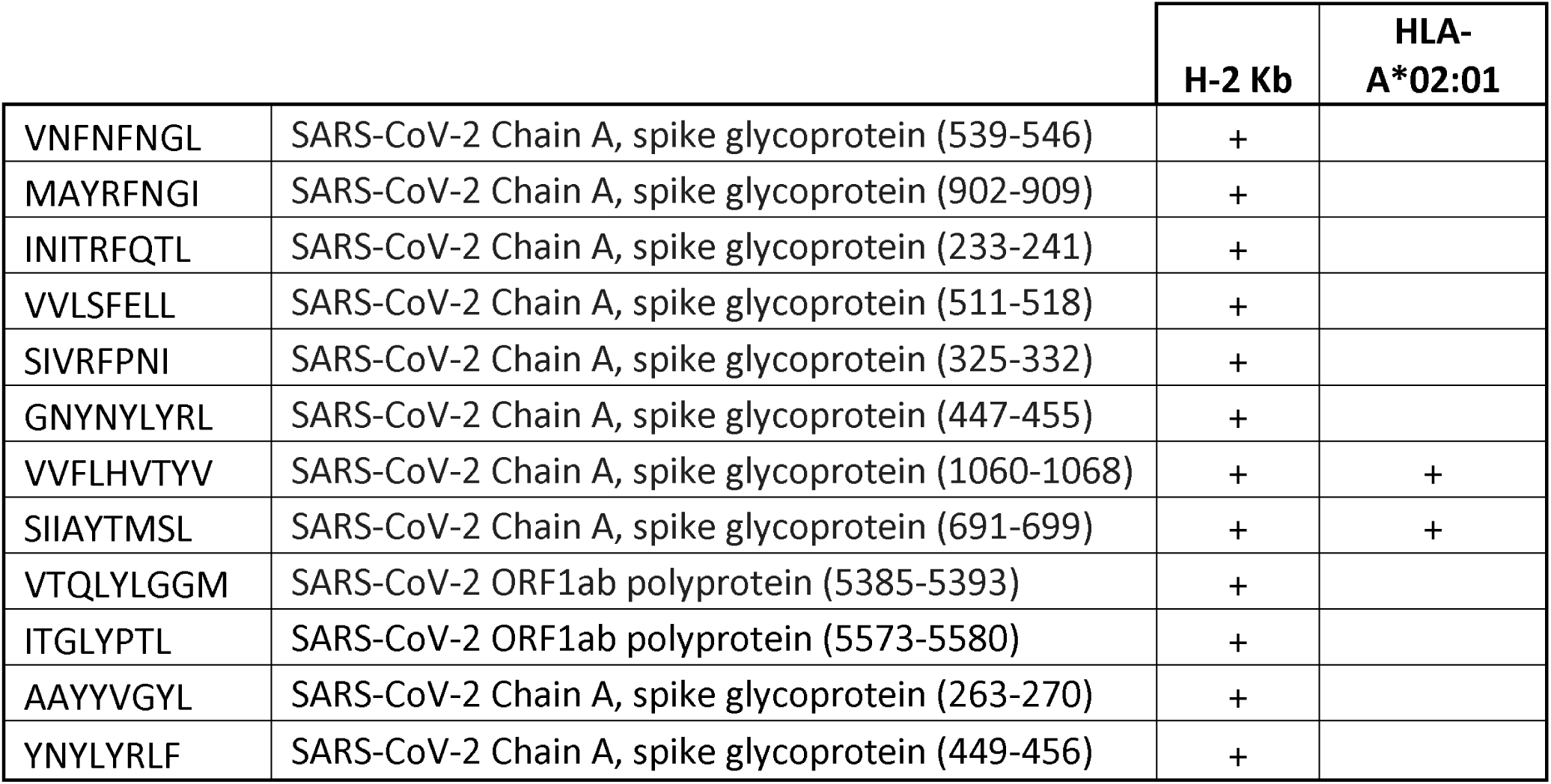
Summary of SARS-CoV-2 peptide binding to mouse H-2 Kb MHC molecules. Peptides from different SARS-CoV-2 proteins were selected for strongest binding on the H-2 Kb mouse allele. After the peptides were found to exchange at least 75% of the exiting QuickSwitch peptide they were tested with the HLA-A2 allele and the cross-reacting peptides were identified.

The exact exchange ratios of the tested HLA-A*02:01 Tetramer-PE molecules with each SARS-CoV-2 peptide can be visualized in Figure 1. All HLA-A2 exchange ratios following a 4-hour incubation can be compared to the exchange ratio of the HLA-A2 Reference Peptide (Figure 1, sample number 29). This peptide has been specifically selected and engineered to have one of the strongest interactions with the peptide binding groove of the HLA-A*02:01 molecule. The Reference Peptide is able to exchange a significant amount of the exiting peptide even when less than 1 µM of it is used for the exchange reaction.

**Figure 1:**
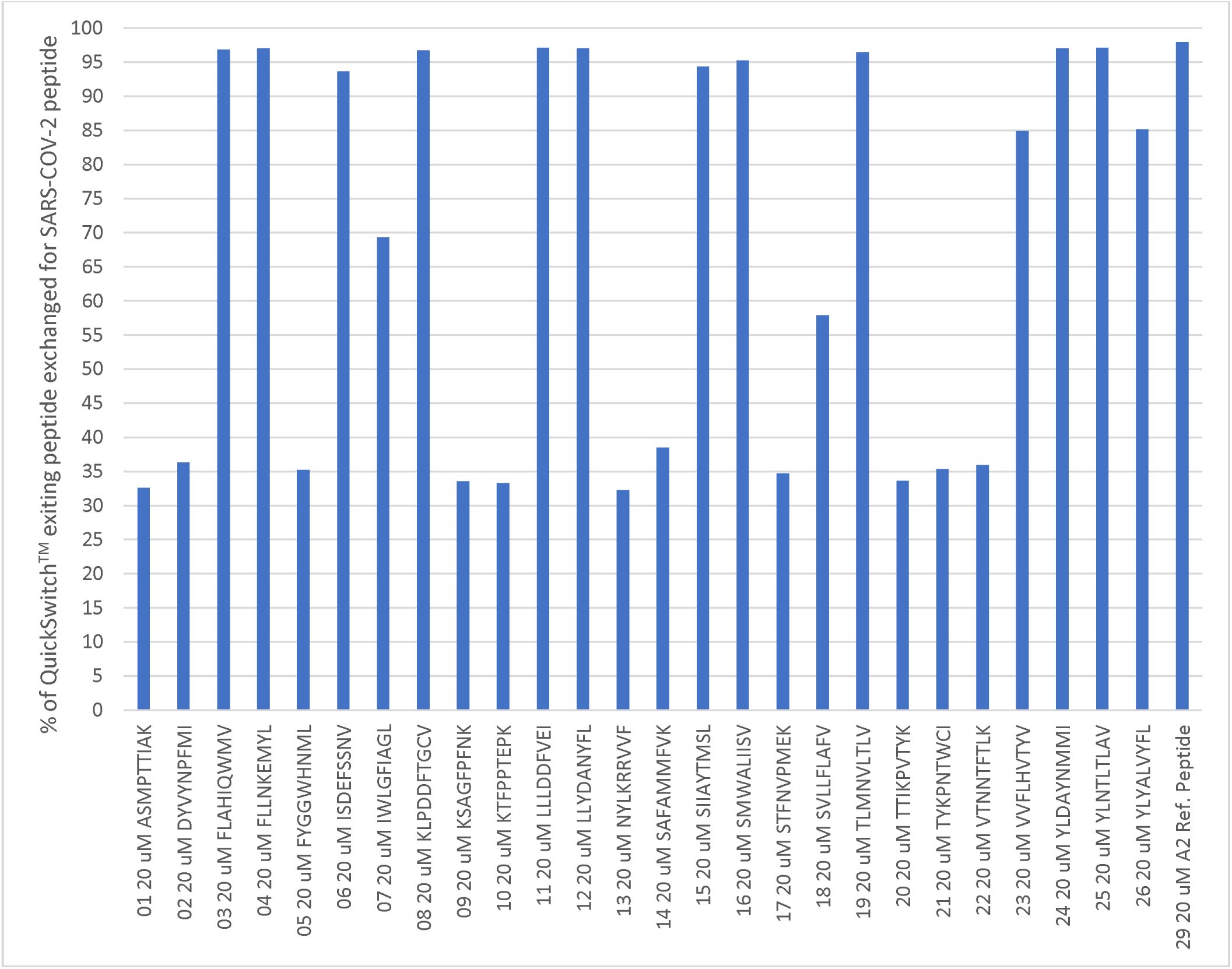
SARS-CoV-2 peptide exchange with QuickSwitch HLA-A*02:01 Tetramer-PE molecules. 50 µg/mL Tetramerized and R-phycoerythrin coupled HLA-A*02:01 molecules were used for the peptide exchange experiment. All HLA-A*02:01 molecules were loaded with a exiting peptide and incubated with 20 µM of competing SARS-CoV-2 peptides for 4 hours at room temperature (≈ 23°C) in the presence of a kit-determined volume of Peptide Exchange Factor #1 (PEF#1). The amount of exiting peptide removed from the MHC molecule was then determined following a capture assay and plotted on the vertical axis for each tested peptide.

As an illustration of the effectiveness of in silico prediction algorithms, we compared the theoretical binding affinity of the studied peptides determined by IEDB and the peptide exchange ratio resulting from the QuickSwitch assay. While most of the peptides determined to have low nanomolar Kd values readily exchange with the exiting peptide of the HLA-A*02:01 molecule, even among the peptides which we have tried there are a few exceptions. Some peptides with low theoretical Kd values show a low binding ability to the QuickSwitch HLA-A2 assay and some peptides that should theoretically be bad binders display a high binding capacity for HLA-A2 (Figure 2). It is those outliers, which may hold the greatest insight into how the immune system may be skewed to resolve an infection or pathology and our QuickSwitch platform provides an effective mechanism to identify those peptides.

**Figure 2:**
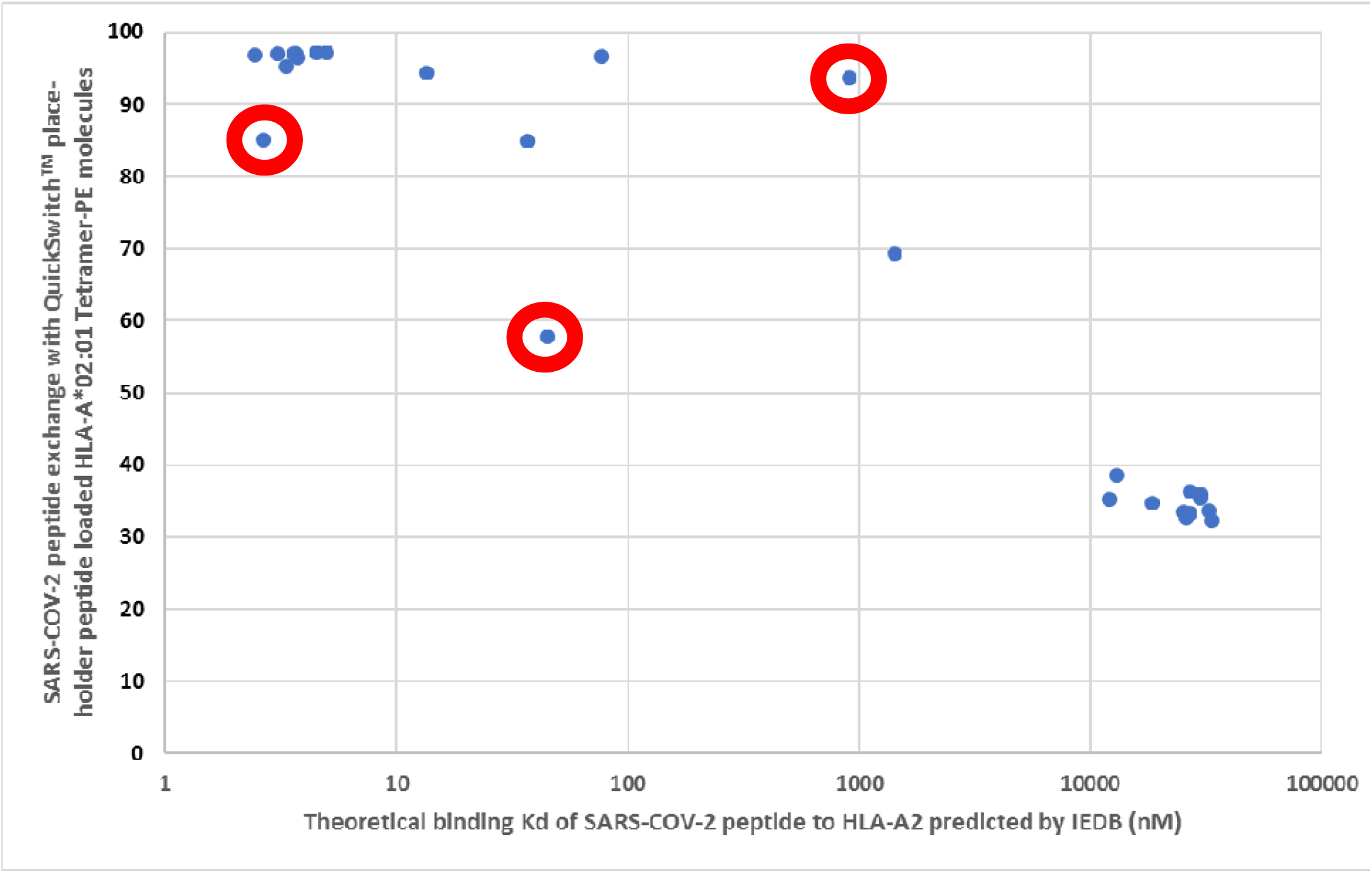
Theoretical binding affinity of SARS-CoV-2 peptides to HLA-A*02:01 compared to the practical binding to a Tetramerized recombinant MHC molecule. Theoretical Kd values of SARS-CoV-2 peptide binding to HLA-A*02:01 molecules as determined by the IEDB web resource were plotted against the resulting HLA-A*02:01 Tetramer-PE peptide exchange values. The peptides outlined in red circles are YLYALVYFL (Theoretical Kd = 2.67 nM, Practical Exchange rate = 85%); SVLLFLAFV (Theoretical Kd = 44.91 nM, Practical Exchange rate = 57.5 %) and ISDEFSSNV (Theoretical Kd = 908.9 nM, Practical Exchange rate = 93.5%).

We tested 19 SARS-CoV-2 peptides selected with the IEDB web resource for their ability to bind to recombinant HLA-DRB1*01:01 (DR1), HLA-DRB1*04:01 (DR4) and HLA-DRB1*15:01 (DR15) molecules. All of the selected peptides were 15 amino acids in length with various binding affinity to each of the three Class II molecules. Unlike with the Class I QuickSwitch platform, the Class II peptide exchange cannot be tested directly on a fluorochrome-conjugated tetramer and the peptide exchange has to be performed on biotinylated recombinant MHC Class II molecules before they are tetramerized following the kit’s guidelines. As a Reference Peptide for all of the HLA-DR molecules tested we used the short sequence of the CLIP_87-101_ peptide – PVSKMRMATPLLMQA. While CLIP is generally believed to be a moderate binder of MHC Class II molecules [11], we have consistently found that its short 15 amino acid form interacts strongly with recombinant DR1, DR4 and DR15 molecules used to make MHC tetramers. Most of the tested peptides bound equally well to all three Class II molecules (Figure 3). Some of the peptide exchanges were repeated with 100 and 10 µM of competing peptide. Peptides RAMPNMLRIMASLVL and SEFSSLPSYAAFATA exchange equally well at 10 µM and 1 mM indicating that they bind strongly to the three alleles (Data not shown). In contrast, peptides IWLGFIAGLIAIVMV, LLLLDRLNQLESKMS and LAFVVFLLVTLAILT did not bind well at the 1 mM level and showed virtually no exchange at 10 µM. HLA-DR molecules seem to have less restrictions to peptide binding than the MHC Class I molecules we have tested. Not all peptides will be equally presented on naturally occurring MHC molecules since some of them will be selected against the antigen processing and presentation system. To fully mimic the naturally occurring peptides, some form of an in vitro biochemical antigen processing model will have to be used to determine which peptides can truly be presented by the antigen presentations system [12–14] and only then should those peptides be tested with the QuickSwitch platform.

**Figure 3:**
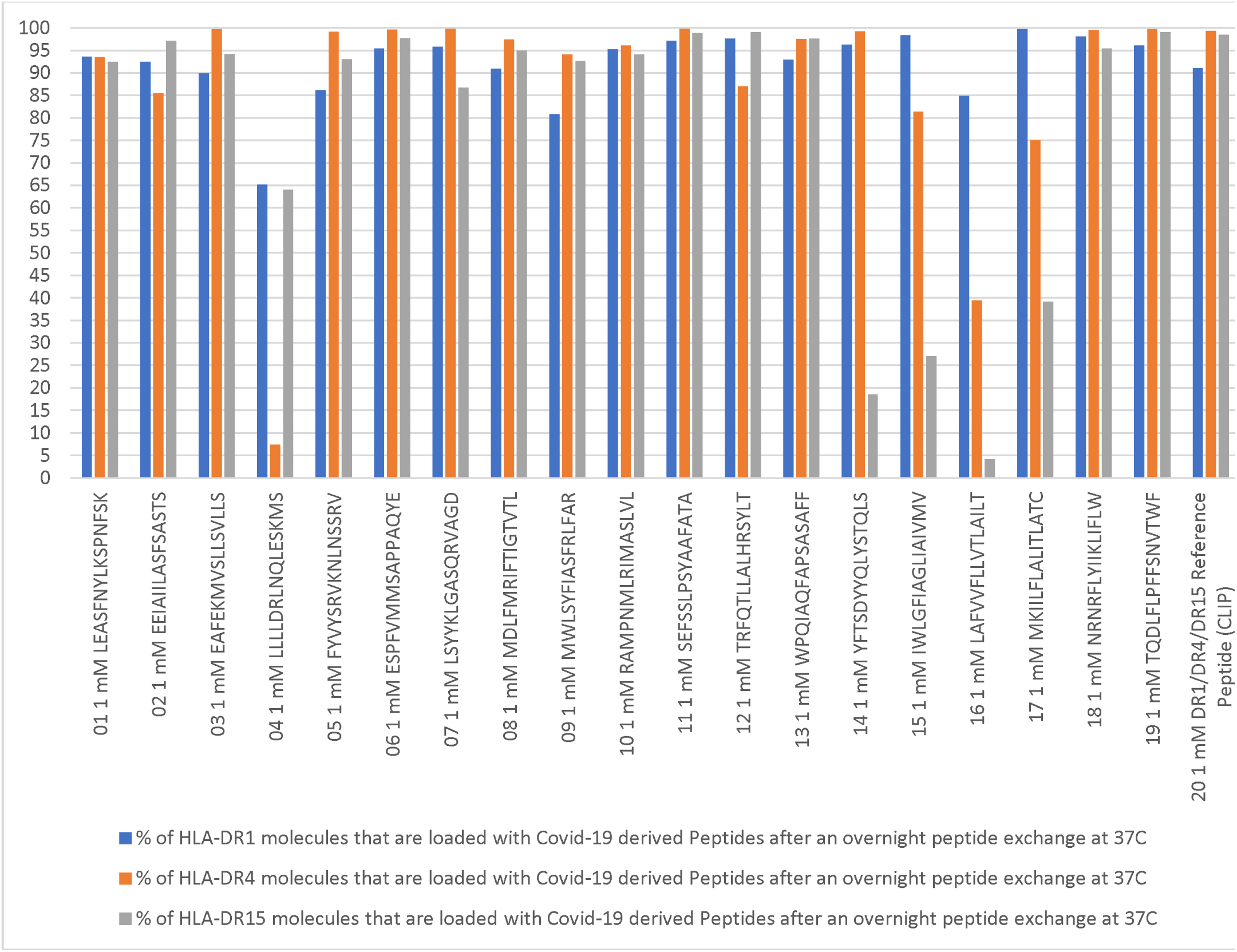
SARS-CoV-2 peptide exchange with QuickSwitch HLA-DRB1*01:01, HLA-DRB1*04:01 and HLA-DRB1*15:01 molecules. 100 µg/mL recombinant HLA-DRB1*01:01 (Blue Bars), HLA-DRB1*04:01 (Orange Bars) and HLA-DRB1*15:01 (Gray Bars) molecules were used for the peptide exchange experiment. All MHC Class II molecules were loaded with an exiting peptide and incubated with 1 mM of competing SARS-CoV-2 peptides overnight (> 10 hours) at 37°C. The amount of exiting peptide removed from the MHC molecule was then determined following a capture assay and plotted on the vertical axis for each tested peptide.

## Discussion

There are concerns that some vaccine researchers focus only on humoral responses to establish the potency and efficacy of their CoV-2 vaccines. This concern stems from the fact that the durability and potency of SARS-CoV-2 vaccines is indeed partially dictated by elicited T cell immune responses. Not examining the CoV-2 T cell immunomonitoring may result in vaccines that induce short lasting humoral responses in the absence of proper T cell memory responses [15–20]. Other than healthy hosts, it is also important to analyze CoV-2-specific T cell responses in certain vaccinated sub-populations such as the elderly, and hosts who bear underlying conditions that are expected to impact the immune system such as diabetes. To better understand the breath of T cell immune responses, three essential steps need to be interrogated, i) it is necessary to identify protein pieces (peptides) that can be processed in antigen processing pathways, ii) to identify those that are able to interact with the groove of MHC molecules, and iii) to validate those peptides with human PBMCs to ensure that there are T cell clonotypes recognizing these peptides in the context of self MHC. This short communication addresses the second item which is MHC occupancy testing of a selected sets of peptides that are reported in the past few months based on either in silica prediction models and/or after validation in PBMCs of hosts. These peptides should be validated in a system other than the one they are predicted in [21]. The QuickSwitch platform for the generation of custom tetramers has proven itself to be a robust tool for the identification of peptides that can form a stable complex with both Class I and Class II MHC molecules. Unlike in-silico systems, the platform used for this study involves the quantitation of real peptide binding to recombinant MHC molecules which can then be used to make tetramers and stain T cells. This allows the users to focus on finding populations of T cells that react to a specific viral peptide without worrying about whether this peptide can be presented by the MHC molecules or if the MHC tetramer that they are using is defective. In addition, the strong-binding peptides that we discovered in this short study can be used for multiple applications and assays. It is expected that these findings can be used by other labs to decipher what viral peptides have the most relevance to fighting the Covid-19 pandemic.

## Acknowledgement

The authors would like to thank Dr. Farshad Guirakhoo for critically reviewing the manuscript.

## Individual contributions

Experiments were conceived by Y.P. and M.C.D. and performed by Y.P. Data analysis was done by Y.P., P.D. and M.C.D. The manuscript was written by Y.P. with editing and contributions from all authors. All authors have given approval to the final version of the manuscript.

